# The scaling of expansive energy under the Red Queen predicts Cope’s Rule

**DOI:** 10.1101/536920

**Authors:** Indrė Žliobaitė, Mikael Fortelius

## Abstract

The Red Queen’s hypothesis portrays evolution as a never-ending competition for expansive energy, where one species’ gain is another species’ loss. The Red Queen is neutral with respect to body size, implying that neither small nor large species have a universal competitive advantage. The maximum population growth in ecology; however, clearly depends on body size – the smaller the species, the shorter the generation length, and the faster it can expand. Here we ask whether, and if so how, the Red Queen’s hypothesis can accommodate a spectrum of body sizes. We theoretically analyse scaling of expansive energy with body mass and demonstrate that in the Red Queen’s zero-sum game for resources, neither small nor large species have a universal evolutionary advantage. We argue that smaller species have an evolutionary advantage only when resources in the environment are not fully occupied, such as after mass extinctions or following key innovations allowing expansion into freed up or previously unoccupied resource space. Under such circumstances, we claim, generation length is the main limiting factor for population growth. When competition for resources is weak, smaller species can indeed expand faster, but to sustain this growth they also need more resources. In the Red Queen’s realm, where resources are fully occupied and the only way for expansion is to outcompete other species, acquisition of expansive energy becomes the limiting factor and small species lose their physiological advantage. A gradual transition from unlimited resources to a zero-sum game offers a direct mechanistic explanation for observed body mass trends in the fossil record, known as Cope’s Rule. When the system is far from the limit of resources and competition is not maximally intense, small species take up ecological space faster. When the system approaches the limits of its carrying capacity and competition tightens, small species lose their evolutionary advantage and we observe a wider range of successful body masses, and, as a result, an increase in the average body mass within lineages.

## 1. Introduction

The widely recognised and extensively debated empirical generalisation known variously as Cope’s Rule, Alroy’s Axiom or even Marsh’s Maxim (Stanley 1973, Polly 1998, Raia & Fortelius 2013) – the tendency for species within a lineage to evolve towards a larger body size, as documented for North American land mammals by Alroy (1998) and for those of the Northern Hemisphere by Huang et al. (2017), does not sit comfortably in modern theoretical frameworks of ecology and evolution. Cope’s basic tenet, that small organisms tend to be unspecialised and increased body size is a result of increased specialisation is difficult to reconcile with general principles of ecology and animal function, such as metabolic scaling or digestive physiology.

Interpreting the empirical patterns that might support Cope’s Rule has also proven fraught with difficulty. Stanley (1973) pointed out, citing Bonner (1968), that small species do not vanish; instead large species become more common. In accord, Jablonski (1997) empirically demonstrated for Cretaceous molluscs that the observed body mass trends arise as the range of body size expands. Raia et al. (2012) found that the later, larger species of Neogene bovids tended to have smaller geographic ranges and shorter geological durations, which they interpreted as support for Cope’s original view of weak directionality of evolution towards increasing specialization. Yet the relationship between specialization and body size is non-trivial and not clearly directional. For example, Pineda-Munoz et al (2016) showed that specialized feeders, such as frugivores, tend to be in the middle rather than the extreme end of body sizes. Collectively, the patterns and analyses attributed to Cope’s Rule suggest an interplay of directional and nondirectional evolutionary processes that requires further explanation.

Body size has often been portrayed as one of the most direct links between microevolution and macroevolution (Maurer et al. 1992, Jablonski 1996). While the diversity of body sizes of organisms has traditionally been explained in terms of microevolutionary processes, macroevolution has been called on to explain selection responsible for body size trends. Stanley (1973) pointed out that explaining Cope’s Rule means explaining why most populations approach optimum body sizes from smaller rather than from larger sizes, a question still largely unanswered. Here we propose that trends in body size appear as a result of a transition from relatively unlimited resource states towards progressively more resource limitation and tighter competition. While it is obvious that infinitely unlimited resource state is never possible, temporarily unlimited resource situations may occur. Mass extinctions can substantially free up previously occupied resources (Raup and Sepkoski 1982, Erwin 2001, Smith et al. 2010), major evolutionary innovations may open resources of new kinds (Heard and Hauser 1995, Raia et al. 2013), and continental drift can open new spaces (Chaboureaua et al. 2014), presenting perhaps the extreme of effectively unlimited resources. The other extreme, characterized by the Red Queen’s hypothesis (Van Valen 1973a, 1980), occurs when resources are occupied to the maximum carrying capacity of the environment, and the only way for species to expand is to outcompete other species via adaptive evolution.

We demonstrate here that under unlimited resources, such as after mass extinctions or after key innovations, small species can expand faster to fill up the available ecological space, and under those circumstances have an evolutionary advantage. When resources in the environment become fully occupied, small species no longer have a universal evolutionary advantage and the range of successful body sizes expands. This happens because under tight competition for resources populations do not grow at the maximum physiologically possible speed, but only at the speed determined by the amount of expansive energy outcompeted from other species. The amount of resources taken away from other species depends on evolving competitive advantages and can be argued to be invariant to body mass (Damuth 1981, 2007).

All else being equal, if resources are not fully occupied, population growth is limited by the generation length, therefore at small body size populations will expand faster (Brown 1995) until the ecological space is filled up, the carrying capacity is reached and competition becomes extremely tight, as assumed by the Red Queen. In such circumstances availability of expansive energy limits population growth (Zliobaite et al. 2017). Therefore, the original Red Queen’s hypothesis describes a the common but also extreme end of the competition spectrum, where body size no has no universal evolutionary advantage.

## 2. How much expansive energy is required for population growth

Van Valen (1980) postulated that natural selection maximises many quantities, but only expansive energy is maximized unconditionally. Trophic energy can only be controlled by live individuals, therefore increasing the control of expansive energy implies population growth. Analysing the scaling of population growth with body mass will clarify how evolutionary competitiveness relates to body size.

### 2.1 Metabolic scaling

The relation between expansive energy and population dynamics is not trivial, due to metabolic scaling. Terrestrial mammals span 24 orders of magnitude from shrews weighing 2 grams to elephants weighing 5 tonnes today, or even 15 tonnes or more in the past Fortelius & Kappelman 1993). Body mass effectively determines the pace of life (Schmidt-Nielsen, 1984). If it was not for metabolic scaling allowing a slower pace of life for larger organisms, all multicellular organisms likely would have to be of one size class. As animals get larger, the surface to volume ratio shrinks, while cumulative transportation distances within the body increase. While energy is required within three-dimensional bodies, acquisition of resources and release of waste happens via surfaces. This dimensionality mismatch between interfaces for energy exchange and units of consumption is the fundamental reason why metabolic rate has to slow down with body size increase.

The dominant metabolic scaling relation, established by Kleiber (1932), is W ∝ M^0.75^, where W is energy consumption rate and M is body mass. Different technical explanations for the scaling exponent have been evoked ranging from supply networks (West et al. 1997, Savage et al. 2008, Brown et al. 2004) to thermodynamics (Ballesteros et al. 2018). Some authors have argued that there is no universal metabolic scaling exponent, only ranges that give the upper and lower bounds (Banavar et al. 2010, Virot et al. 2017). Even though the exact numeric exponent is still debated (White and Seymour 2003), it is clear that the metabolic rate cannot scale isometrically. This makes a fundamental difference in energy allocation: large organisms are relatively more efficient in maintenance, but slower in reproduction (Brown 1995).

### 2.2 Scaling of population densities

While the metabolic rate and thus energy consumed by an individual depends on body size, energy consumption of the whole population of a species is expected to be invariant to body size (Van Valen 1976). Damuth (2007) demonstrated, using simulation models, that when randomly chosen species evolved to take energy from other species, population densities settle at an inverse of the metabolic rate.

This scaling of population densities is fundamentally related to the number of individuals that can be supported by a constant amount of energy E ∝ M^0^. If one individual consumes energy at rate W, then the number of individuals that can be supported with that energy scales as E/W ∝ M^0^/M^0.75^ = M^−0.75^. Indeed, empirical evidence suggests (Damuth 1981, Marquet 2002, Jetz et al. 2004) that globally population density of primary consumers scales as N ∝ M^−0.75^, which is the inverse of the metabolic scaling.

### 2.3 Population growth rate when resources are unlimited

Ecological theory and observations suggest that the maximum population growth rate scales as R_growth_ ∝ M^−0.25^ (Fenchel 1974, Brown 1995). The negative exponent implies that, given enough resources, small species can expand their populations faster. From the definition of the growth rate (R_growth_ = N_new_/N) it follows that the increase in the number of individuals scales as N_new_ = R_growth_ N ∝ M^−0.25^ M^−0.75^ = M^−1^, where N is the number of individuals the previous time step. The expansive energy -- that is, the extra energy needed to support those new individuals per unit time scales as E_expansive_ = W N_new_ ∝ M^0.75^ M^−1^=M^−0.25^.

This means that in order to sustain population growth at the physiologically possible maximum speed, populations of smaller species need to acquire more expansive energy per unit time. This can work for relatively short time intervals, when the sources from which the extra energy comes are effectively unlimited. Sooner or later, however, the carrying capacity of the environment is reached and the main means to acquire expansive energy becomes competition with other species. In such circumstances small species would need to compete harder in order to sustain their maximum population growth rates.

### 2.4 The Red Queen’s population growth rate (when resources are maximally occupied)

Damuth (2007) argued that, in the Red Queen’s domain, the bulk amount of resources to be taken from competitors at the population level depends on the evolutionary advantage but does not dependent on the body mass of individuals. We will analyse this condition later in this paper; assuming for now that it holds, we can derive the population growth rate in the Red Queen’s domain.

If acquired expansive energy at the population level does not depend on body mass (E_expansive_ ∝ M^0^), then the number of new individuals this energy can support scales as N_new_ = E_expansive_ /W ∝ M^0^/M^0.75^ = M^−0.75^. The number of individuals at the previous time step scales the same as the population density N ∝ M^−0.75^. The population growth rate therefore scales as R_growth_ = N_new_/N ∝ M^−0.75^/M^−0.75^ = M^0^, and is invariant to body size of individuals.

The remainder of the study analyses demographic and evolutionary implications of this scaling.

## 3. Demographic paths to evolutionary expansion

Population growth rate is a statistical construct, which we decompose to understand the mechanics of its scaling. We analyse in which ways expansive energy can affect population growth, and possible resultant constraints.

### 3.1 Scaling of life history parameters

Birth and mortality rates, as most life history descriptors, including aging and longevity, closely relate to the metabolic scaling (Lindstedt and Calder 1981, Brown et al. 2004, Hulbert et al. 2007). Smaller animals grow faster, reproduce and die sooner; per lifetime animals are expected to endure the same number of heartbeats, breaths or chews (Peters 1983, Fortelius 1985), independently of their body mass. Life span is finite primarily because of intrinsic wear of the organism due to metabolic process (Fortelius 1989, Savage et al. 2006, Hulbert et al. 2007). Since the speed of life, the metabolic rate per unit of body mass, scales as M^−0.25^ and the durability of an organism (how many heartbeats, breaths, chews) scales as M^0^, the maximum lifetime theoretically should scale as the durability divided by the speed of use, ∝ M^0^/M^−0.25^ = M^0.25^.

While maximum lifespan defines a physiological lifetime, ecological longevity, described as life expectancy at birth, captures both extrinsic and intrinsic causes of death. Life expectancy is an average over all the individuals in the population, including those that die shortly after birth. Scaling literature for animals often reports either one or the other, but both are expected and reported to scale with body mass with the same theoretical exponent of 0.25 (Polishchuk et al. 1999). The ecological life expectancy determines how many offspring can be produced at the population level.

Life expectancy does not directly contribute to population growth, but mortality rates do. Under a coarse assumption of uniform mortality rate across the age groups^1^, however, scaling of mortality rates directly follows from ecological life expectancy, one scaling as the inverse of the other (Keyfitz and Caswell 2005). Since life expectancy theoretically scales as ∝ M^0.25^, mortality rate should scale as R_death_ ∝ M^−0.25^. The empirical mortality rate of mammals has been reported to scale very closely to this, with the exponent of −0.24 (McCoy and Gillooly 2008).

When a population does not grow, the birth rate scales with the same exponent as the mortality rate R_birth_ = R_death_ ∝ M^−0.25^. Indeed, annual fecundity^2^ of mammals has been observed to scale with the exponent of −0.26 (Hamilton et al. 2011). Since life expectancy scales with the exponent of 0.25, and the reproductive lifetime scales with the same exponent, the expected total number of offspring per lifetime is independent of body mass, which was first pointed out by Williams (1966). This is a logical and the only possible scaling for a population that does not to grow.

### 3.2 Ways of population growth

A population grows if birth rates exceed mortality rates. For example, if the birth rate per population per year is 25% and the death rate is 20%, the population will grow 5% per year. This relationship follows from the definition of growth rate R_growth_ = N_new_ / N = (N_born_ - N_died_) / N = N_born_ / N - N_dead_ / N = R_birth_ - R_death_, where N is the number of individuals at the previous time step, N_born_ is the number of individuals born during this period, N_dead_ is the number of individuals that died during this period, and R_birth_ and R_death_ are the birth and mortality rates correspondingly.

This relationship suggests that there are two ways for expansive energy to make population grow – either to increase the birth rate, or reduce the mortality rate. Increase in the birth rate can happen in two ways: physiological or demographic. A demographic increase would mean a change in population structure such that the fraction of reproductive females increases, and can only happen if the population is shrinking, which is not relevant for this analysis. A physiological increase would mean more frequent birth or more offspring per birth per reproductive female. While specialized life history strategies are widely recognized (Sibly and Brown 2009), birth rates are governed by metabolic scaling (Lindstedt and Calder 1981) and are not easy to change permanently without physiological tradeoffs. Elevated reproductive rates lead to shortened lifespan, as limited resources enforce a compromise between investment in reproduction and somatic maintenance (Edward and Chapman 2011, Blacher et al 2017).

More likely, availability of expansive energy due to an evolutionary advantage would increase average life expectancy, which is to a large extent extrinsic. While the maximum lifespan is physiological, and by and large cannot be altered by making life more comfortable (West 2017), the average lifespan, since it is ecology and competition driven, can be so altered. Therefore, increased life expectancy does not mean extending the life span of the oldest individuals, but adding more time throughout all age classes. Such effects have been observed by Tidiere et al. (2012), who analysed mortality rates of zoo and wild animals and showed that being at a zoo reduces the mortality rates of all age groups by a roughly constant factor. Lowering mortality rates below equilibrium would make a population grow.

An evolutionary advantage that can help to acquire energy more effectively – for instance, by walking faster or shorter distances to the next food, higher probability of finding food, higher chewing or digestion efficiency -- can reduce mortality rates by reducing the risk of exposure to predators, getting into an accident, or getting sick. For instance, Van Valkenburgh (2009) demonstrated that the risk of tooth fracture for predators increases when competition is tighter. While eliminating one cause of death may slightly increase probabilities to die from other causes (Keyfitz 1977), for clarity of exposition we assume in this analysis that all possible causes of death for a species fall under a compound probability and the evolutionary advantage changes that probability.

### 3.3 Effects on mortality rates with unlimited resources

Let the exact current population size be N = aM^−0.75^ (scaling reviewed in Section 2.4 holds, since N ∝ M^−0.75^). Let the exact birth and mortality rates of a non-growing population be R_birth_ = R_death_ = bM^−0.25^. The number of individuals that would die per period is N_died_ = aM^−0.25^ bM^−0.75^ = abM^−1^.

Suppose an unlimited amount of resources suddenly becomes available and the population starts growing at a maximum physiological rate. As demonstrated earlier, expansive energy required to sustain such growth scales as E_expansive_ ∝ M^−0.25^. Since the individual metabolic rate scales as ∝ M^0.75^, the number of individuals that can be supported by this energy scales as ∝ M^−0.25^M^−0.75 =^ M^−1^. Denote the exact number of individuals that escaped death due to availability of extra energy as N_saved_ = cM^−1^. Then the adjusted mortality rate due to expansive energy is R_growth_ * = (abM^−1^ - cM^−1^)/aM^−0.75^ = bM^−0.25^ - (c/a)M^−0.25^ = R_death_ - (c/a)M^−0.25^.

Assuming that the evolutionary advantage proceeds in steps and has a small effect on mortality rates at a time, such that the population age structure stays the same, the population growth rate is given by R_growth_ * = R_birth_ - R_death_ * = R_birth_ - (R_death_ - (c/a)M^−0.25^) = (c/a)M^−0.25^. The coefficients a and c do not depend on body mass, thus, population growth rate scales as ∝ M^−0.25^. Therefore, when resources are effectively unlimited smaller species have a competitive advantage for expansion.

### 3.4 Effects on mortality rates in the Red Queen’s realm

In the Red Queen’s domain expansive energy becomes available only if competitors lose their share of energy. Suppose one species acquires an evolutionary advantage that allows it to acquire extra energy. Further assume, in line with Damuth (2007), that the amount of this energy at the population level depends on the magnitude of the evolutionary advantage, but not on the body size of individuals, such that E_expansive_ ∝ M^0^.

Since the individual metabolic rate scales as ∝ M^0.75^, the number of individuals that can be supported by this energy scales as ∝ M^−0.75^. Denote the number of individuals saved from death due to the acquired expansive energy at the population level as N_saved_ = dM^−0.75^. Then, the adjusted mortality due to expansive energy is R_death_ * = (abM^−1^ - dM^−0.75^) / aM^−0.75^ = bM^−0.25^ - d/a = R_death_ - d/a.

Assuming that the effect on mortality rates is small, such that the population age structure and the birth rates stay the same, the population growth rate is given by R_growth_* = R_birth_ - R_death_* = R_birth_ - R_death_ + d/a = d/a. The coefficient d/a relates to the amount of expansive energy acquired and does not depend on body mass. Thus, in the Red Queen’s domain, population growth rate does not depend on body size, given that the evolutionary advantage is equally beneficial for all body sizes. The latter condition remains to be analysed.

## 4. How the Red Queen’s competitive evolution can be neutral with respect to body mass

Our remaining question is under what conditions and in what circumstances an individual evolutionary advantage can be such that the expansive energy at the population level would depend only on the magnitude of that advantage, but not on body size.

### 4.1 Limits on species expansion

Suppose a species acquires a small evolutionary advantage; for example, an extra crest on their teeth. We assume that the probability of an acquisition of a new trait for an individual does not depend on body size, that is, a mouse and an elephant are equally likely to acquire a given trait. The reality is, of course, more complicated as understood by population genetics (Loewe and Hill 2010). At the genomic level various interdependencies are likely, including different lengths of genome and the fact that the same traits may be encoded differently in different organisms. Yet we argue that from the biomechanical perspective as a coarse approximation the assumption is realistic. For example, a hypocone – a cusp that extended the occlusal surface of the upper molar teeth -- convergently evolved more than 20 times among mammals during the Cenozoic (Hunter and Jernvall 1995).

Within the scope of this analysis we assume that the new trait gives a competitive advantage in energy acquisition. This can manifest itself as walking faster or requiring shorter distances to reach the next food item, higher probability of finding food, higher chewing or digestion efficiency, or faster escape from predators, allowing longer or more concentrated periods of feeding. Not all morphological traits would fall under this assumption. For example, an increase in dental crown height would extend the durability of teeth, but would not directly increase effectiveness of day-to-day resource acquisition. We assume that the evolutionary advantage is such that it does not change the body mass, does not extend lifespan, does not directly give more offspring, and that the individual metabolic rate stays the same.

As argued earlier, we assume that an advantage in energy acquisition reduces mortality rates throughout all age groups, by allowing, for instance, better predator avoidance, less risk taking while foraging, and staying healthier due to quality nourishment. We assume, for simplicity and in the spirit of species being separately evolving units (Simpson 1961), that every individual within the species shares the same trait. More precisely, we assume that within-species variation is less than between species variation, as is typically the case for functionally important traits (Polly et al. 2017).

We model the Red Queen’s competition for energy as a proportional prize contest (Cason et al. 2018), in which rewards are shared in proportion to performance. The winning species do not gain unlimited resources, the gain is only proportional to the competitive advantage they have acquired. While eventually “new adversaries grinningly replace the losers” (Van Valen 1973), within an arbitrarily short time period losing species co-exist and control a share of resources. This is conceptually different from a winner-takes-all contest (Cason et al. 2018), which in our analysis would correspond to the scenario of unlimited resources. This latter kind corresponds to allometric scaling of population growth as seen in the previous sections – small species expand exponentially faster.

In the Red Queen’s realm, a competitive advantage can only give a tiny gain in expansive energy at a time. The proportional prize contest implies that the gain in expansive energy is relative to the other species’ collective performance. Thus, the availability of expansive energy, rather than the generation length, is the limiting factor. Even if smaller species have a shorter generation length, they cannot utilize their reproductive advantage at a full capacity unless they can manage to be more efficient in energy acquisition using the same tools. In the hypocone example this would require a cusp of the same shape to provide relatively more chewing capacity for small mammals than for large mammals. If the same trait leads to the same biomechanical effects, energy gain rather than the generation length would be the limiting factor for species expansion, and body size would not have a universal evolutionary advantage.

The question is how biomechanical advantages balance out with different metabolic rates. Even if the biomechanics of the same trait are assumed to be independent of body size, food intake does depend on body size and scales allometrically with it (Shipley et al. 1994). It remaining to show how this balances out at the population level.

### 4.2 The Red Queen’s competitive evolution as a proportional prize contest

Our analysis assumes a closed environment of a finite carrying capacity, which remains fixed during the period of analysis. For simplicity of exposition, assume that there are only two species and that at the initial time step the system is at equilibrium, such that neither species is expanding or declining.

We conceptualize the proportional prize competition with the following model. Energy is renewable. At every time period (e.g. a day), a constant amount of energy is generated in the environment. Assume that energy comes in a standardized form, everybody can extract the same amount of energy from a unit of food, and the same time of day is available for eating for everybody. At every period (e.g. every day), everybody starts eating and is eating until food available on that day runs out. There is not enough food for complete satisfaction, just enough for survival.

Let W_1_ be the metabolic rate of species A, and W_2_ be the metabolic rate of species B, neither of which will change during the period of analysis. We model the efficiency of resource acquisition as the average speed of food acquisition throughout the period. This does not necessarily mean instantaneous speed up of intake (eating faster), but rather that that less time per day spent on foraging (finding food faster).

Let us define the speed of food acquisition at the equilibrium to be proportional to the metabolic rate S_1_^(0)^ ∝ W_1_, and S_2_^(0)^ ∝ W_2_. The superscript (0) denotes the starting time at which populations are at equilibrium. Let the number of individuals of the two species at the equilibrium be N_1_ and N_2_. Let T denote the fraction of day that both species spend on foraging. Then the trophic energy controlled by each population is E_1_^(0)^ = N_1_S_1_^(0)^T^(0)^, and E_2_^(0)^ = N_2_S_2_^(0)^T^(0)^. The carrying capacity of the environment is then the sum of energies controlled by the two populations K = E_1_^(0)^ + E_2_^(0)^.

Assume further that the first species evolves a favourable trait which, for the time being, makes acquisition of resources more efficient. Denote this evolutionary advantage as α > 1, such that the speed of food acquisition becomes S_1_ = αS_1_^(0)^. Assume that the second species meanwhile does not evolve any new traits, and thus its speed of eating remains the same, that is, S_2_ = S_2_^(0)^.

Since species A has evolved an advantage and can now acquire energy faster, the total amount of energy that can be acquired per individual during the next period changes. Individuals of species A become slightly better nourished, while individuals of species B become slightly more undernourished. We hypothesize that this primarily affects mortality rates, as analysed in the previous section, and changes in mortality rates lead to population growth or decline. Analytically, that happens in the following way.

When species A gets an evolutionary advantage, the overall foraging time for both species gets shorter. The equilibrium state provides just enough food for everybody, but when species A speeds up food acquisition, the daily carrying capacity of the environment is exhausted faster. The foraging time for both species thus becomes T = βT^(0)^, where T^(0)^ is the daily foraging time for both species at the equilibrium, and β < 1 is a coefficient that depends on the initial population sizes of both species. From the assumption that the carrying capacity of the environment stays constant, we can find the expression for β. The carrying capacity is equal to the amount of energy both species control before and after acquiring the evolutionary advantage E_1_^(0)^ + E_2_^(0)^ = E_1_ + E_2_.

Assuming that the number of individuals does not change during the first step of analysis, the amount of energy obtained by species A during the period is E_1_ = N_1_S_1_T = N_1_ αS_1_^(0)^βT^(0)^ = αβE_1_^(0)^.

The amount of energy obtained by the second species is E_2_ = N_2_S_2_T = N_2_S_2_^(0)^βT^(0)^ = βE_2_^(0)^.

From these equations we can express the foraging coefficient as β = (E_1_^(0)^ + E_2_^(0)^) / (αE_1_^(0)^ + E_2_^(0)^).

Denote the initial proportion of trophic energy controlled by species A as π = E_1_^(0)^/K. Then, β = 1 / (απ + 1 − π). Since species A now can obtain energy faster, the population will control energy in excess of the equilibrium metabolic rate. This can be thought as the extra energy that improves the probability of survival and this way allows the population to grow, as analysed in the previous section. It remains to analyse how the population growth rate depends on the evolutionary advantage α.

The amount of energy obtained by species A can at equilibrium metabolic rates support N_1_* individuals in the next period, where N_1_* = E_1_/(W_1_T^(0)^) = N_1_αS_1_^(0)^βT^(0)^/(W_1_T^(0)^) ∝ N_1_αW_1_^(0)^β/W_1_= N_1_αβ. Since the growth rate is defined as the change in the number of individuals, as compared to the previous number of individuals R_expansion_ = (N_1_*−N_1_)/N_1_ = N_1_*/N_1_ − 1 = N_1_αβ/ N_1_ − 1 = αβ − 1, where α is the coefficient denoting the evolutionary advantage of species A, and β is the coefficient related to the share of the carrying capacity initially held by species A.

Plugging in the expression for β obtained earlier gives the population growth rate for species A at the first time step after acquiring the evolutionary advantage R_expansion_ ∝ αβ − 1 = α / (απ + 1 − π) − 1.

Similarly, the rate of decline for species B is given by R_decline_ ∝ β − 1 = 1 / (απ + 1 − π) − 1.

From these expressions we see that the rates of expansion and decline only depend on how much of trophic energy species control at the beginning, and on the evolutionary advantage α. If, as we argue, α does not depend on body mass, the rest does not depend on body mass, from which follows a mechanistic explanation for macroevolution rates under the Red Queen – the expansion and decline rates do not depend on body size.

### 4.3 Patterns of continued expansion and decline

We have analysed the scaling of expansion and decline over one time step, which can be of an arbitrary length and during which we can reasonably assume no change in the population size. In practice, of course, evolutionary advantages improve gradually and expansions as well as declines may take many steps. Interestingly, over many steps the expansion resulting from an evolutionary improvement saturates, even if no counter evolutionary advance follows from other species. This links to the unimodal patterns of species’ rise and decline observed in the fossil record and living clades (Jernvall and Fortelius 2004, Foote et al. 2007, Liow and Stenseth 2007, Tietje and Kiessling 2013, Lim and Marshall 2017).

These unimodal patterns, known as hat patterns, are fully compatible with the Red Queen’s hypothesis (Zliobaite et al. 2017), prescribing maximum competition. In the previous work we attributed unimodality at continued evolution to inertia – a memory-like process that presumably due to constraints on adaptation due to building upon already existing characteristics carries species expansion or decline forward over multiple time steps. Indeed, if a species acquires an evolutionary trait that gives a relative advantage in energy acquisition, the population starts to grow. Intuition may suggest that once the evolutionary advantage has been acquired, the population growth will continue at the same rate for many time steps, at least until another species acquires a corresponding evolutionary advantage, but this is not the case.

Consider the following stochastic simulation of species expansion over multiple time steps. Suppose we have one species happily exploiting a fixed share of resources in some constant environment. Let some of the individuals acquire an evolutionary advantage that improves the efficiency of foraging. Assume that about 1% of the population acquires the advantage and branches off to make a new species. The simulation will show how under limited carrying capacity of the environment this acquisition of an advantage gradually leads to extinction of the old type. This simulation is not the same as our earlier theoretical analysis. The simulation takes our earlier theoretical analysis forward to illustrate by an example a multiple time steps scenario.

To align with our analysis in the previous sections let us call the species with the evolutionary advantage species A, and the remaining species without an advantage – species B. At the start the population size of species B is much larger than that of the branching-off species A. After every time step, population sizes rebalance to match the expansive energy captured by each during that period.

Figure 1 depicts how such populations would change over time. Since no more evolution in traits happens during the analysis period, all the observed growth and decline is due intrinsic inertia following the first evolutionary advantage.

**Figure 1.**
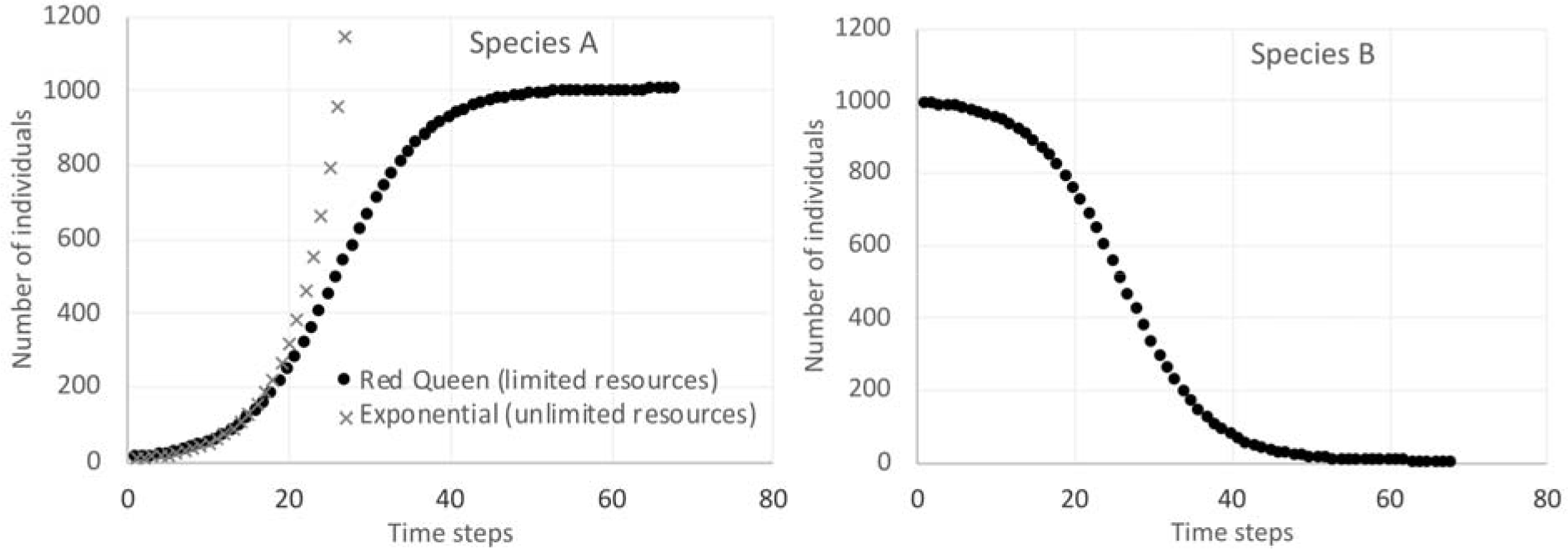
Simulation of continued growth with a fixed evolutionary advantage. Species A has an evolutionary advantage over species B at time 0. Simulation parameters: α=1.2, K=1000, W_1_=W_2_=1, N_A_^(0)^=10, N_B_^(0)^=990.

The grey trajectory in the figure demonstrates exponential growth, which holds if effectively unlimited resources are available. This is how the growth in species abundance, occupancy or range would look if the evolutionary advantage would be a major key innovation, leading to, for instance, opening of new major resources.

In the Red Queen’s domain, resources are utilized to full capacity and the only way to advance in expansion is to outcompete others. Here we see that the advantaged species first expands rapidly, but after a while expansion saturates, even if no other species come up with a counter-matching evolutionary advantage. Species B declines following the same pattern but in reverse.

Even though the carrying capacity is fixed, at the beginning the population growth looks indistinguishable from exponential. Since the population of species A is very small at the beginning, as their expansive energy comes from species B and the population of species B is so large, this makes the amount of resources effectively unlimited. Yet, the magnitude of an evolutionary advantage determines the end of expansion (approaching the peak of the curve). As the population of species A grows larger, competition against other species turns into within-species competition and saturates the growth. Such logistic growth patterns are well established in ecological modelling under limited carrying capacity (Verhulst 1838, Pearl and Reed 1920). For decline to occur, competition from rapidly evolving (and therefore expanding) daughter species has been invoked as the “seed of decline” hypothesis (Fortelius et al. 2014).

The shape of the expansion curve and the time of saturation depends on the initial populations’ sizes, body sizes and the magnitude of the evolutionary advantage. This intrinsic perspective on saturation contributes to the understanding of the hat-like patterns in the fossil record (Jernvall and Fortelius 2004, Foote et al. 2007, Liow and Stenseth 2007, Tietje and Kiessling 2013).

## 5. The Red Queen offers an explanation for Cope’s Rule

Our study portrays a simplified version of evolutionary and demographic processes, yet for the first time it links together seemingly contradictory patterns – the directional trend of increasing body mass over time, known as Cope’s Rule, and the Red Queen’s hypothesis, portraying species expansion and decline as a stochastic process.

The Red Queen’s competition depicts the extreme end of evolutionary circumstances, which is also expected to be the most common. In the Red Queen’s world, resources are always fully occupied, therefore, all else being equal, differential growth of competing species, should not occur without decline of other species. The other extreme, only possible theoretically, is a world with resources so plentiful that anyone who wants gets plenty and there is never a need to compete for resources. In reality, resources can become nearly unlimited for short times, either through environmental change, extinction, or evolution unlocking new resources. Under partially occupied resources, species compete to some extent, but one species’ gain does not necessarily translate to an equivalent loss for all other species, which allows a faster expanding species to outcompete a slower one.

This continuous perspective towards competition for resources offers an elegant explanation for the trends of average body mass increase observed within lineages the fossil record. When the system is far from the limit of resources, small species can expand faster and take the ecological space. Then when the system approaches the limit of the carrying capacity and competition tightens, large and small species become equally competitive, which explains the expanding range of sizes towards the larger end of the spectrum and thus the increase in mean size. The gradual transition from unlimited resources to a zero-sum game for resources explains Cope’s Rule.

Our results have implications for interpreting mass extinctions. It has been considered that small body size is favoured for survival during mass extinctions, the effect is known as the “Lilliput Effect” (Harries and Knorr 2009, Twitchett 2007) An alternative hypothesis might be that mass extinctions affect all body sizes uniformly, but small body size is instead favoured during the initial recovery, as, for instance, evidence of marine vertebrates suggests (Sallan and Galimberti 2015).

The concept of fast and slow life strategies is well established in ecology, as the r/K selection theory (MacArthur and Wilson 1971, Pianka 1970, Reznik 2002, Brown et al. 2004). The theory postulates that r-selected organisms have short life spans, are small and quick to mature. They typically live in unstable, unpredictable environments, where the ability to reproduce rapidly is important. K-selected organisms occupy more stable environments. They are larger in size, have longer life expectancies, are stronger or are better protected and generally are more energy efficient (Cunningham et al. 2001). Our new perspective towards competition for resources at different intensities directly links this ecological theory with observed macroevolutionary trends.

## Acknowledgements

We thank Stein Vidar Haugan for an insightful discussion following our blog post that raised curious questions and provided inspiration for this analysis. Research leading to these results was partially supported by The Academy of Finland grant to IŽ (no. 314803). This is a contribution from the Valio Armas Korvenkontio Unit of Dental Anatomy in Relation to Evolutionary Theory.

## Footnotes

1 This assumes that the mortality rate is constant across the age groups. In reality young and old individuals of a natural population are generally more likely to die than middle aged individuals (Tidiere et al. 2016), but it is a useful approximation commonly taken to allow generic demographic analysis (Keyfitz and Caswell 2005).

2 Annual fecundity is reported per female, while the birth rate is defined per population, but since the number of reproducing females is proportional to the total population size (Polishchuk et al. 1999), these two rates are bound to have the same scaling exponent.

